# Sex and gonadectomy modify behavioral seizure susceptibility and mortality in a repeated low-dose kainic acid systemic injection paradigm in mice

**DOI:** 10.1101/2023.05.22.541824

**Authors:** Niraj V. Lawande, Elisabeth A. Conklin, Catherine A. Christian-Hinman

## Abstract

**Objective:** Sex differences in epilepsy appear driven in part due to effects of gonadal steroids, with varying results in experimental models based on species, strain, and method of seizure induction. Furthermore, removing a main source of these steroids via gonadectomy may impact seizure characteristics differently in males and females. Repeated low-dose kainic acid (RLDKA) systemic injection paradigms were recently shown to reliably induce status epilepticus (SE) and hippocampal histopathology in C57BL/6J mice. Here, we investigated whether seizure susceptibility in a RLDKA injection protocol exhibits a sex difference, and whether gonadectomy differentially influences response to this seizure induction paradigm in males and females.

**Methods:** Adult C57BL/6J mice were left gonad-intact as controls or gonadectomized (females: ovariectomized, OVX; males: orchidectomized, ORX). At least 2 weeks later, KA was injected i.p. every 30 minutes at 7.5 mg/kg or less until the animal reached SE, defined by at least 5 generalized seizures (GS, Racine stage 3 or higher). Parameters of susceptibility to GS induction, SE development, and mortality rates were quantified.

**Results:** No differences in seizure susceptibility or mortality were observed between control males and control females. ORX males exhibited increased susceptibility and reduced latency to both GS and SE, but OVX females exhibited increased susceptibility and reduced latency to SE only. However, ORX males, but not OVX females, exhibited strongly increased seizure-induced mortality.

**Significance:** The RLDKA protocol is notable for its efficacy in inducing SE and seizure-induced histopathology in C57BL/6J mice, the background for many transgenic strains in current use in epilepsy research. The present results indicate that this protocol may be beneficial for investigating the effects of gonadal hormone replacement on seizure susceptibility, mortality, and seizure-induced histopathology, and that gonadectomy unmasks sex differences in susceptibility to seizures and mortality not observed in gonad-intact controls.

## 1. Introduction

Multiple aspects of epilepsy and seizure susceptibility exhibit sex differences.^1,2^ For example, in humans the epilepsies as a whole are slightly more common in males,^3–5^ but the degree and presence of a sex difference typically varies with the specific etiology and type of epilepsy.^1^ In animal models, several studies have documented sex differences in acute seizure susceptibility or chronic epilepsy presentation, although the directionality of the sex effect can vary by model and species.^6–14^ Many of these sex differences are likely a result of distinct neural effects of gonadal hormones in males and females.^1,15–17^ Both estrogen and progesterone have proconvulsant and anticonvulsant effects, as documented in studies examining different species and models of seizure and epilepsy induction.^21–35^ Testosterone can also have mixed effects on seizures due to metabolism producing both anticonvulsant and proconvulsant compounds.^36^

In previous studies, removal of testes in male rats (orchidectomy, ORX) resulted in decreased susceptibility to seizure induction by kainic acid (KA) and bicuculline.^15,37,38^ However, ORX male rats also exhibited increased susceptibility to picrotoxin-induced generalized seizures (GS), whereas ORX male mice had no change in susceptibility to pentylenetetrazole-induced seizures.^39,40^ In contrast to the mixed effects of ORX on seizures, removal of ovaries in females (ovariectomy, OVX) is documented to increase seizure susceptibility in both rats and mice in response to various chemoconvulsants.^28,37,40–46^ By contrast, another study found increased severity of systemic KA-induced behavioral seizures in ovary-intact female rats.^38^ Overall, these results indicate that gonadectomy can have differing effects on seizures between males and females, with effects likely shaped by the specific mode of seizure induction and the experimental species.

Recent reports describe the advantages of repeated low-dose KA (RLDKA) administration in inducing status epilepticus (SE) and histopathology characteristic of temporal lobe epilepsy in C57BL/6J mice,^47–49^ which can be more resistant than other strains to a single bolus injection of KA^50–52^ and are the background for many transgenic strains in current use in epilepsy research. The RLDKA paradigm is also highly adaptable, as the timing and dosage of drug administration can be modified in accordance with differences in mouse strain or other factors,^48,49,53^ potentially providing more experimental control over the severity of induced seizures than is achieved with single bolus administration of a high KA dose.^50–52^ However, previous studies using RLDKA injections only examined males or did not differentiate data collected from males vs. females,^47–49,54^ leaving open the question as to whether sex differences are observed in response to RLDKA seizure induction. In addition, determining effects of gonadectomy would clarify the utility of this model for studies examining the effects of gonadal hormone replacement and the selective roles of neural sources of these hormones in modulating seizure susceptibility and excitability. Therefore, here we tested the susceptibility to RLDKA induction of behavioral GS and SE, and seizure-induced mortality, in gonad-intact and gonadectomized male and female C57BL/6J mice.

## 2. Methods

### 2.1 Animals

All animal procedures were approved by the Institutional Animal Care and Use Committee of the University of Illinois Urbana-Champaign. C57BL/6J mice (n = 33 females, n = 25 males) were purchased from Jackson Laboratories for delivery at 6 weeks of age. All mice were group-housed (2-5 per cage) and were given a minimum of 4 days to acclimate to our animal facility prior to experimental handling. Mice were housed on a 14:10 h light:dark cycle with food (Teklad 2918, an irradiated diet) and water available *ad libitum* and lights-off at 1900 h.

### 2.2 Gonadectomy surgeries

19 females underwent OVX and 10 males underwent ORX. Additionally, 3 females and 3 males were included as shams. For OVX/ORX procedures, mice were anesthetized by isoflurane inhalation (Clipper Distributing Company, St. Joseph, Missouri). Prior to surgery, mice were given the analgesic carprofen (5 mg/kg s.c., Zoetis). Bilateral OVX was performed as previously described.^55^ For ORX, a small midline incision was made in the skin on the ventral side of the scrotum. The tunica was pierced and each testis was gently exposed and removed by cautery. Bupivacaine (0.25%, 7 ul, Hospira, Lake Forest, Illinois) was applied locally into each surgical site for postoperative analgesia.^55^ Surgical sites were closed with wound clips, which were removed 7-10 days later. All OVX/ORX mice were allowed 2 weeks for recovery from the procedure and to allow for full depletion of circulating gonadal hormones before seizure induction testing (described below). Successful OVX status was confirmed by visual inspection of the shrunken uterus following euthanasia.^56^ Sham surgeries were conducted identically but without removal of the ovaries and without tunica incision and removal of the testes.

### 2.3 Repeated low-dose kainic acid injection

Age-matched OVX/ORX, sham, and gonad-intact control mice (9-11 weeks of age) were tested in a RLDKA paradigm adapted from a previous report.^49^ Injections began between 1100-1600 h. KA (Tocris Bioscience #0222) was dissolved in sterile 0.9% saline at 2 mg/ml and stored at - 20°C for up to 7 days prior to use. On the day of injection, the mice were brought to the experiment room, weighed, individually placed in fresh cages, and given 30 minutes to acclimate. KA was i.p.-injected at 7.5 mg/kg every 30 minutes until the animal developed SE, defined by at least 5 GS of Racine stage of 3 or higher using the following definitions: stage 3 – wet dog shakes, head nodding, and unilateral forelimb clonus; stage 4 – stage 3 with bilateral forelimb clonus or rearing; stage 5 – stage 4 with repeated rearing and falling or tonic-clonic seizures.^57^ An additional dose was administered if the animal did not have a subsequent GS for at least 30 minutes after initial seizure onset. One OVX female received only one injection because stage 5 seizures presented at 30 min after injection. No further injections were given once a cumulative seizure burden of 30 (calculated by the sum of all Racine scores ≥ 3) was reached. Six OVX and one control females with particularly severe seizures that were nearing this cumulative seizure burden score were given half doses of KA, and the data were adjusted accordingly. 2 hours after the last injection, all mice were injected s.c. with 500 μl Lactated Ringer ‘s solution or 0.9% saline and returned to the mouse facility in single cages. Mortality was quantified based on survival to 300 min from the first injection. At 3 or 7 days after injection, mice were deeply anesthetized with pentobarbital (110 mg/kg i.p.) and euthanized by intracardiac perfusion with 4% paraformaldehyde.

### 2.4 Statistical analyses

Statistical comparisons were done using R or OriginPro (OriginLab, Northampton, MA). Data from sham mice were highly similar to the data collected from the naïve gonad-intact mice (i.e., those that did not undergo sham surgery) of the corresponding sex (**Figure S1**). Therefore, the sham mice were combined with the intact mice as the control male and female groups in analyses.

Data normality was determined using Shapiro-Wilks tests. In the absence of a reliable non-parametric equivalent of a two-way ANOVA, data transformations (log or square root) were used where specified to allow for use of parametric statistics. Choice of log vs. square root transformation was based on accuracy of the normalizing effect for the given dataset. To test for differences in the number of injections needed to produce GS and SE, and the latency in minutes to first GS or SE, two-way ANOVA and Tukey ‘s *post hoc* tests were used with sex and gonad status as the independent variables. For latency measurements, the time of first injection was designated as time 0. Fisher ‘s exact tests and *post hoc* Bonferroni-corrected pairwise comparisons were used to test for differences in the proportions of mice in each group developing GS or SE. Kaplan-Meier analysis with log rank tests for equality of survival times was used to compare differences in latency to death between the groups. A power analysis demonstrated that the final sample sizes achieved statistical power for 2-way ANOVA over 84% at α = 0.05 for a medium effect size (Cohen ‘s f) of 0.3. Data are presented as means ± SD. Statistical significance was set at p < 0.05.

## Results

### 3.1 Body weight is increased in gonadectomized female mice but not males

Gonadectomized mice can exhibit changes in body weight in comparison to age-matched gonad-intact controls, often resulting in increased body weight in OVX females and decreased weight in ORX males.^58–61^ To determine whether this effect was observed by the time of seizure testing in this cohort, we analyzed the body weight measured on the day of RLDKA injection. Body weight differed significantly by sex (F_1,54_ = 71.72, p < 0.001) and gonad status (F_1,54_ = 6.73, p = 0.012), with a significant interaction between sex and gonad status (F_1,54_ = 13.72, p < 0.001) (**Figure 1**). Pairwise comparisons indicated that, as expected, control males (n = 15, 24.71 ± 2.15 g) had higher body weight than control females (n = 14, 19.64 ± 1.17 g) (p < 0.001). This sex effect persisted when comparing ORX males (n = 10, 24.25 ± 1.69 g) to OVX females (n = 19, 22.26 ± 1.08 g) (p = 0.011). However, whereas OVX females exhibited higher body weight than control females (p < 0.001), ORX and control males were not different (p = 1). These results indicate that OVX females, but not ORX males, exhibited increased weight gain within 2-4 weeks after gonadectomy.

**Figure 1.**
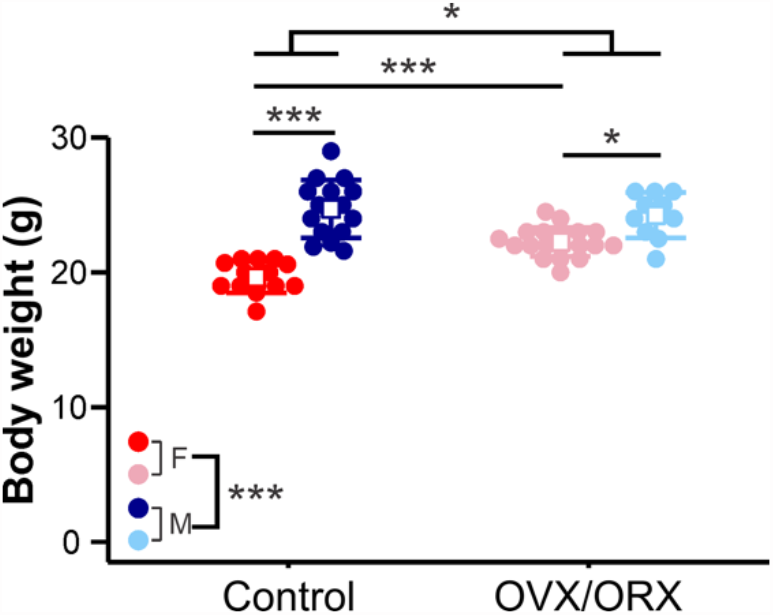
Body weight is increased at 2-4 weeks after gonadectomy in females but not males. Individual values (circles) and mean ± SD (squares and lines, respectively) for body weight measured on the day of RLDKA injection. Red = control females (F); blue = control males (M); pink = OVX females; light blue = ORX males. *p<0.05, ***p<0.001

### 3.2 Gonadectomy increases susceptibility to GS in males

To determine whether GS induction in response to RLDKA injection is influenced by mouse sex and/or gonad status, male (control n = 15, ORX n = 10) and female (control n = 14, OVX n= 19) mice were tested. The data for the number of injections needed to produce GS and latency to the first GS were log- and square root-transformed, respectively, to account for non-normal distributions. The number of injections varied significantly by gonad status (F_1,54_ = 20.544, p < 0.001) but not by sex (F_1,54_ = 0.041, p = 0.841) with no interaction between sex and gonad status (F_1,54_ = 1.38, p = 0.245) (**Figure 2A**). Fewer injections were required to induce GS in ORX males compared to controls (controls 3.67 ± 0.82, ORX 2.30 ± 0.48 injections, p = 0.0022), and there was a trend towards significance between control and OVX females (controls 3.50 ± 1.29, OVX 2.65 ± 0.79 injections, p = 0.061). There were no sex differences observed within control (p = 0.89) or gonadectomized groups (p = 0.778).

**Figure 2.**
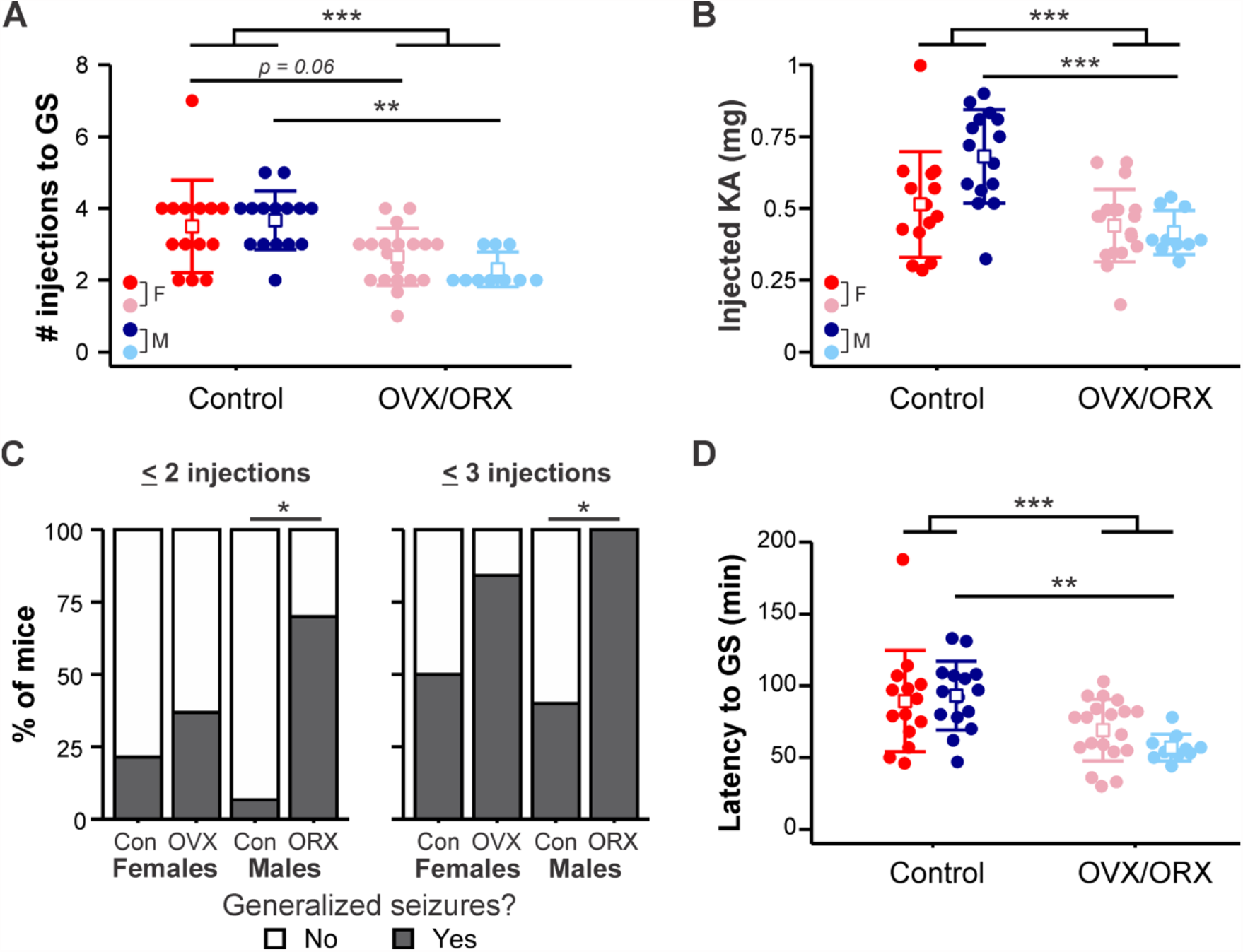
Effect of gonadectomy on susceptibility to GS is most prominent in males. A) Individual values (circles) and mean ± SD (squares and lines, respectively) for the number of i.p. kainic acid injections needed to reach GS (GS). Raw data are presented; for statistical analysis, the data were log-transformed to improve normality. B) Individual values and mean ± SD for the total amount of KA injected to initiate GS. C) Percentage of mice in each group with at least one GS by two (left) or three (right) KA injections. D) Individual values and mean ± SD for the latency in minutes to reach GS. Raw data are presented; for statistical analysis, the data were square root-transformed to improve normality. Red = control females (F); blue = control males (M); pink = OVX females; light blue =ORXmales.^*^p<0.05,^**^p<0.01,***p<0.001

Given the differences in body weight observed between the groups (**Figure 1**) and that dosing was normalized to body weight, we calculated the total amount of injected KA required to induce GS. The amount of KA differed by gonad status (F_1,54_ = 18.56, p < 0.001) and there was an interaction between gonad status and sex (F_1,54_ = 5.95, p = 0.018), with no effect of sex on its own (F_1,54_ = 3.32, p = 0.074) (**Figure 2B**). More KA was required to induce GS in control males (0.68 ± 0.16 mg) than in ORX males (0.42 ± 0.08 mg, p < 0.001), but there was no difference between control females (0.51 ± 0.18 mg) and OVX females (0.44 ± 0.13 mg, p = 0.95).

There were also differences between groups in the proportion of mice that had at least one GS by 2 injections (p = 0.005) and by 3 injections (p = 0.002) (**Figure 2C**). A greater proportion of ORX males than control males had at least one GS within both 2 injections (control, n = 1 of 15, ORX n = 7 of 10; p = 0.010) and 3 injections (control, n = 6 of 15, ORX, n = 10 of 10; p = 0.017). No differences were found in rate of GS development between OVX and control female groups at either 2 injections (controls n = 3 of 14, OVX n = 7 of 19; p = 1) or 3 injections (controls n = 7 of 14, OVX n = 16 of 19; p = 0.34).

Similarly, there was an effect of gonad status (F_1,54_ = 15.488, p < 0.001; **Figure 2D**) on the latency to the first GS but no effect of sex (F_1,54_ = 0.151, p = 0.699) and no interaction between sex and gonad status (F_1,54_ = 1.917, p = 0.172). Furthermore, the effect of gonadectomy was primarily driven by the difference within the male group, as ORX males exhibited shorter latency to the first GS than corresponding controls (controls 93.13 ± 23.92, ORX 56.90 ± 9.33 min, p = 0.0049). In females, however, there was no difference between OVX and control groups (controls 88.00 ± 37.45, OVX 69.11 ± 21.42 min, p = 0.219). These results indicate that gonadectomy increased susceptibility to GS induction in response to RLDKA injection in males.

### 3.3 Gonadectomy increases SE susceptibility in both males and females

We also determined whether sex and/or gonadectomy impact susceptibility to SE induction. Four mice (1 from each group) did not survive to SE and thus were not included in these analyses. The number of KA injections needed to reach SE differed by gonad status (F_1,50_ = 29.452, p < 0.001) but not by sex (F_1,50_ = 1.184, p = 0.282) and with no interaction between sex and gonad status (F_1,50_ = 0.079, p = 0.779) (**Figure 3A**). As with susceptibility to initial GS, fewer injections of KA were required to induce SE in ORX males compared to controls (controls 4.14 ± 0.66, ORX 2.67 ± 0.71 injections, p = 0.0025). Furthermore, OVX females also displayed an increased susceptibility to SE induction (controls 4.35 ± 1.07, OVX 3.02 ± 1.05 injections, p = 0.0013). The data for the amount of KA required to induce SE were log-transformed to account for non-normal distributions. There was an effect of gonad status (F_1,50_ = 22.43, p < 0.001) with no effect of sex (F_1,50_ = 1.36, p = 0.25) and no interaction (F_1,50_ = 1.62, p = 0.21) (**Figure 3B**). More KA was required to induce SE in control males (0.77 ± 0.13 mg) than ORX males (0.48 ± 0.13 mg, p < 0.001), but there was only a trend towards significance for the comparison between OVX (0.50 ± 0.17 mg) and control females (0.63 ± 0.15 mg, p = 0.061). In addition, the proportions of mice that reached SE by 2 (p = 0.002) or 3 (p < 0.001) injections varied significantly between groups (**Figure 3C**). Although no two groups differed significantly in the rate of SE induction within 2 injections by pairwise testing, a larger proportion of ORX males (n = 8 of 9) reached SE by 3 injections compared to both control males (n = 2 of 14) (p = 0.004) and control females (n = 2 of 13) (p = 0.009), with a trend approaching significance in comparison to OVX females (n = 11 of 18) (p = 0.069). The rates in OVX and control female groups were not different (p > 0.15).

**Figure 3.**
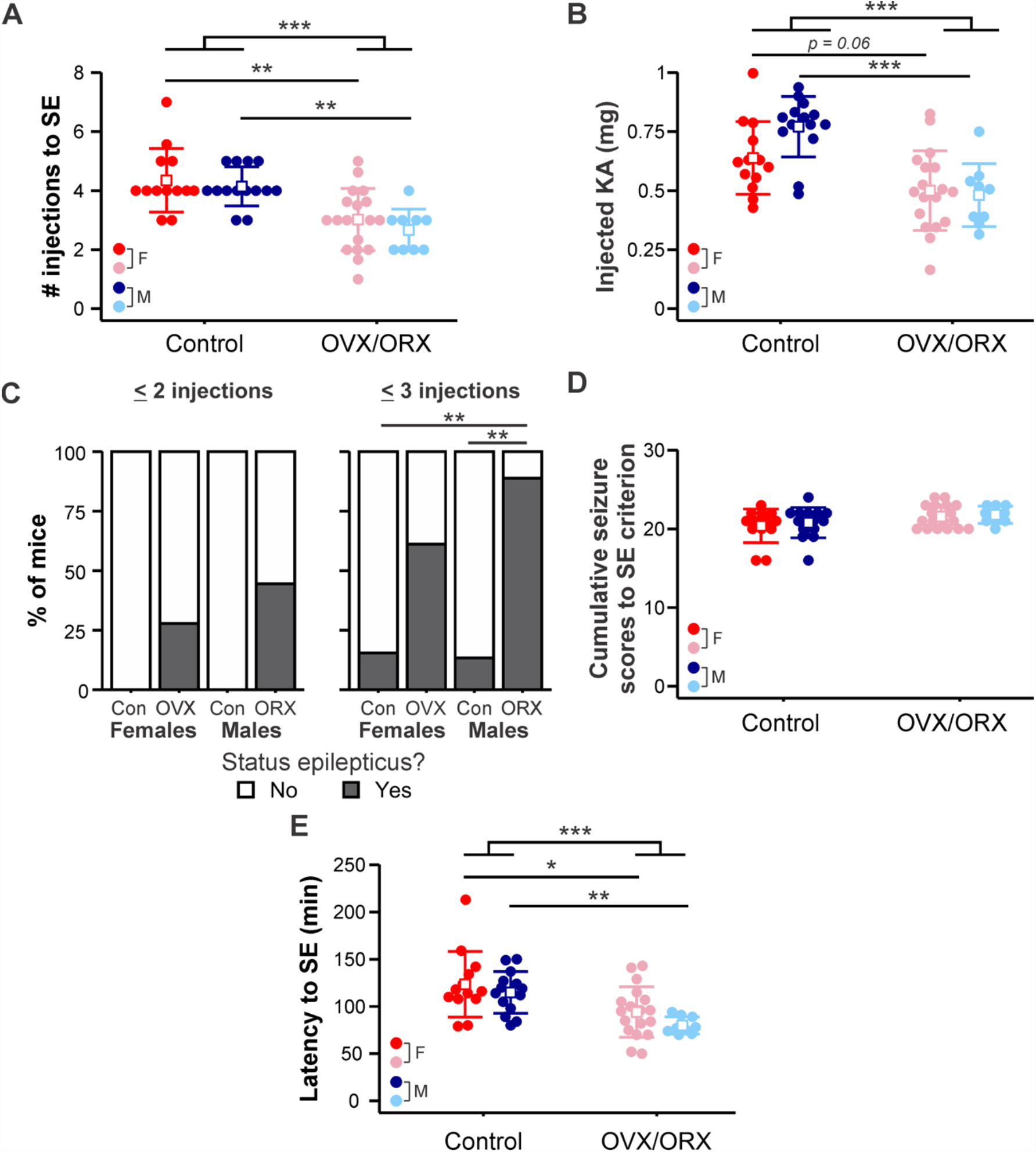
OVX females and ORX males exhibit increased susceptibility to status epilepticus (SE). A) Individual values (circles) and mean ± SD (squares and lines, respectively) for the number of kainic acid injections needed to reach SE. B) Individual values and mean ± SD for the total amount of KA injected to initiate SE. Raw data are presented; for statistical analysis, the data were log-transformed to improve normality. C) Percentage of mice in each group that developed SE by two (left) or three (right) KA injections. D) Individual values and mean ± SD for the cumulative Racine scores of the first 5 GS (i.e., to SE criterion). Raw data are presented; for statistical analysis, the data were log-transformed to improve normality. E) Individual values and mean ± SD for the latency in minutes to reach SE. Raw data are presented; for statistical analysis, the data were square root-transformed to improve normality. Red = control females (F); blue = control males (M); pink = OVX females; light blue = ORX males. ^*^ p<0.05, ^**^p<0.01, ^***^p<0.001

To further assess SE susceptibility, we quantified the cumulative Racine score seizure burden of the first 5 GS (Racine stage 3 or higher) to assess the severity of seizures leading up to SE criterion. These data were log-transformed to account for non-normal distributions. There were no effects of sex (F_1,50_ = 0.44, p = 0.51) and no interaction between sex and gonad status (F_1,50_ = 0.04, p = 0.85) (**Figure 3D**). Although there was a moderate effect of gonad status in two-way ANOVA (F_1,50_ = 5.03, p = 0.029), *post hoc* pairwise comparisons did not indicate a significant effect of OVX in females (p = 0.25) nor OVX in males (p = 0.54), indicating that the severity of seizures leading up to SE was not affected by sex or gonadectomy.

The data for latency in minutes to SE were square root-transformed to account for non-normal distributions. Gonad status (F_1,50_ = 20.854, p < 0.001), but not sex (F_1,50_ = 2.236, p = 0.141), was found to have a significant effect on the latency to SE, with no interaction (F_1,50_ = 0.23, p = 0.63) (**Figure 3E**). Gonadectomized mice of both sexes exhibited shorter latency to SE than corresponding controls (males: controls 114.86 ± 22.17, ORX 79.89 ± 9.03 min, p = 0.0089; females controls 123.46 ± 34.68, OVX 92.67 ± 24.62 min, p = 0.014). These results indicate that susceptibility to SE in response to RLDKA injection was increased following gonadectomy in both male and female mice.

### 3.4 Gonadectomy increases susceptibility to seizure-induced mortality in males but not females

Although previous studies indicate high survival rates following RLDKA injection,^48,49^ not all mice in the present cohort survived the injection regimen and the induced SE. To determine if this mortality reflected differences based on sex and/or gonad status, we conducted a Kaplan-Meier analysis examining 300 min after the first KA injection to encompass the time of the injection series and the immediate 2-hour post-injection observation period. ORX males (n = 7 of 10 died) exhibited significantly shorter survival times compared to control males (n = 1 of 15 died, p < 0.001) and OVX females (n = 2 of 19 died, p = 0.0017) (**Figure 4**). Control females (n = 4 of 14 died) were not different from either OVX females (p > 0.23) or control males (p > 0.13). When only OVX mice with full doses were included in analyses, the difference between OVX females and ORX males persisted (p = 0.018) and the lack of difference between OVX and control females remained. These results indicate that ORX males, but not OVX females, exhibit greatly increased susceptibility to mortality in response to this RLDKA paradigm.

**Figure 4.**
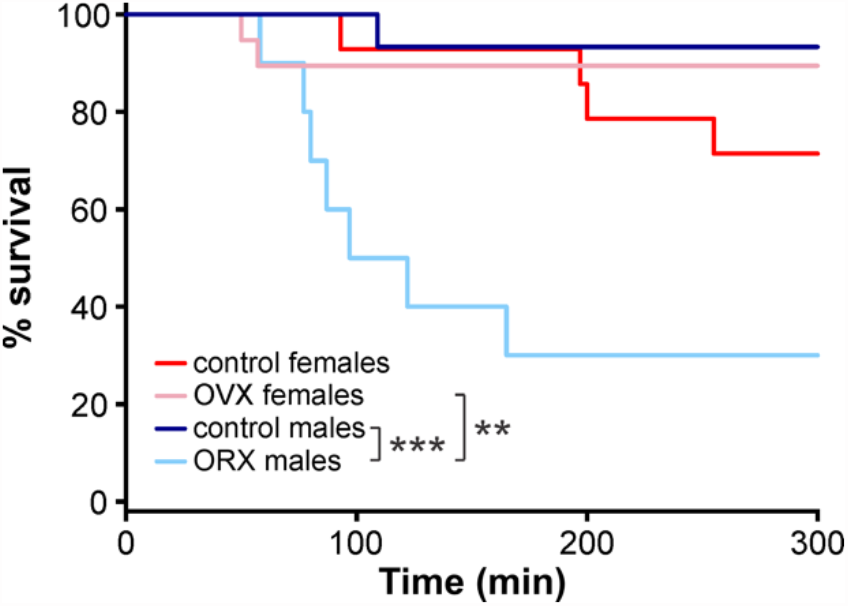
ORX males exhibit increased seizure-related mortality. Kaplan-Meier plot for percent survival over 300 min starting from the time of the first KA injection. ^**^p<0.01, ^***^p<0.001

## Discussion

RLDKA paradigms were recently found to be highly effective in inducing seizures, SE, and post-SE histopathology in C57BL/6J mice.^47–49^ However, it is unclear whether mouse sex and/or gonad status influence susceptibility to seizures and SE in this model. Such information would indicate whether this model may be useful in studies of sex differences and effects of gonadal hormones in seizure activity. The present results demonstrate that gonadectomy had a detrimental effect, driving increased susceptibility to GS and SE. With respect to susceptibility to GS, the effect was largest in males, whereas both ORX males and OVX females exhibited increased susceptibility to SE. In addition, ORX males, but not OVX females, exhibited increased seizure-related mortality, and the rate of mortality was higher in ORX males than OVX females. These studies thus indicate that gonadectomy increases sensitivity to seizure and SE induction in this model, with more prominent effects in males, and reveals sex differences not observed in the gonad-intact condition.

In females, the depletion of ovarian progesterone may drive the increased susceptibility to SE after OVX. Progesterone primarily exerts anticonvulsant effects through its metabolite allopregnanolone, which potentiates GABA_A_receptor-mediated synaptic inhibition.^22–25,29,35,62,63^ However, progesterone actions via nuclear progesterone receptors can also increase excitatory firing and exacerbate seizures.^27,32,33^ Removal of the ovaries also depletes circulating ovarian estradiol. Although estradiol has largely been found to have proconvulsant effects, it is important to note that anticonvulsant effects have also been reported depending on the dose, receptors activated, brain region, and whether acute or chronic seizures were studied.^21,24,26,28,30,31,38,64–67^ Therefore, reduction of estradiol may at least partially underlie the observed seizure-promoting effects of OVX in the present studies, which are in line with several previous results.^28,37,40–45^ The present findings indicate that the RLDKA paradigm may be useful in studies examining the effects of physiological replacement of circulating progesterone and/or estradiol on acute seizure and SE susceptibility, and in studies examining the brain itself as a source of these hormones.^38,68^

Although the findings of increased mortality in ORX male mice would indicate an overall reduced seizure threshold, it should be noted that previous results on the effect on seizures of ORX in male rodents have been mixed. One study documented increased susceptibility to picrotoxin-induced GS in ORX male rats,^39^ in line with the present results in ORX mice. By contrast, others found the reverse effect on KA-induced seizures, with ORX rats exhibiting a decrease in GS, and increased latency to seizures of both low and high severity.^15,38^ Of note, rats and mice exhibit some species differences in responses to seizure-inducing drugs, including KA.^69,70^ One possible explanation for these discrepant results could be differential rates of metabolism of testosterone to estradiol vs. androgens^36,71^ in different species or strains. Alternatively, differences between the present study and past work may reflect differences in the mode of KA administration. While the present work tested a systemic RLDKA paradigm, the other studies that contrast with these results used single bolus injections of larger doses of KA.^15,38^ Follow-up studies directly comparing susceptibility to RLDKA, single KA, and other chemoconvulsants in ORX vs. testis-intact animals would help to clarify the degree to which these different results reflect variations in strain, species, methodology, or other factors.

In line with previous findings using RLDKA paradigms in C57BL/6J mice,^48,49^ the present results demonstrate high survival rates for control males and females. However, although both control and OVX females had mortality rates comparable to intact males, ORX males had a strikingly higher mortality rate, with 70% dying within the acute time frame of the injection series. As a result, using this RLDKA protocol to study post-SE sequelae in ORX males will likely require protocol modifications such as reduced dose or increased interval between injections. Conversely, the increased rate of mortality in ORX C57BL/6J males may suggest that comparison of testis-intact and ORX males in this RLDKA paradigm may be useful in identifying mechanisms associated with sudden unexpected death in epilepsy.^72,73^

The finding of higher mortality in ORX males suggests that testis-derived factors provide critical neuroprotection in male mice, and indicates that sex steroids made outside of the gonads (e.g. in the brain itself) may yield greater compensation for the loss of gonadal steroids in females than in males. Indeed, gonadectomy can differentially impact levels of sex hormones, and the corresponding synthesizing enzymes, in males compared to females.^74–76^ For example, gonadectomy decreases estradiol levels in the preoptic area of male, but not female, rats.^76^ In females, ovarian hormones produce prominent neuroprotective effects.^77–80^ In this regard, it is important to note that in the present study, six OVX females were given decreased doses of KA at later time points in the injection series due to early expression of severe seizures, which may have provided some degree of protection against mortality. However, the amount of KA administered to induce SE was not different between gonadectomized males and females, despite the higher body weight of ORX males. Therefore, increased mortality in ORX males cannot be ascribed to a difference in the amount of KA administered.

Several studies indicate differences in acute seizure susceptibility in female rodents at different stages of the estrous cycle.^1,78–80^ The present experiments, however, were not designed nor powered to examine effects of estrous cycle stage. Therefore, testing the RLDKA paradigm at different stages of the estrous cycle would be an interesting follow-up study. Of note, no sex differences in seizure or SE susceptibility were observed in controls. Therefore, in many studies using this model, inclusion of mixed groups of males and females should be highly feasible, thus meeting requirements of granting agencies for incorporation of both male and female animals in experiments.^84^

Previous research in C57BL/6J mice indicates that RLDKA effectively induces seizure-associated neuropathology and gliosis,^47–49^ which are often lacking in this strain when seizures are induced by a single peripheral injection of a higher dose of KA.^50–52^ Given the present differences in seizure susceptibility, neuropathological assessments in control and OVX/ORX mice following RLDKA injection are pertinent future studies.

A limitation of the present work is that seizure evaluation was done by screening for behavioral seizures without electroencephalography (EEG). It thus remains possible that more subtle effects of sex and gonadectomy on seizures, particularly early on in the injection series, might be observed. These studies indicate that follow-up work using EEG and examining the effects of selective gonadal hormone replacement in the RLDKA paradigm would be fruitful avenues of future investigation.

## Author contributions

C.A.C.-H. designed research; N.V.L. and E.A.C. performed research; E.A.C., N.V.L., and C.A.C.-H. analyzed data; C.A.C.-H., E.A.C., and N.V.L. wrote the paper.

## Acknowledgements

We thank Karen Wilcox and Ashley Zachery-Savalla for helpful discussions on implementing the repeated low-dose kainic acid injection protocol. This work was supported by the National Institutes of Health (NIH)/National Institute on Aging through grant R21 AG077694 (C.A.C.-H.) and a Jenner Family Research Fellowship from the School of Molecular and Cellular Biology at the University of Illinois (N.V.L.).

## Conflicts of Interest

The authors have no conflicts of interest to disclose.

## Supporting Information

**Figure S1.**
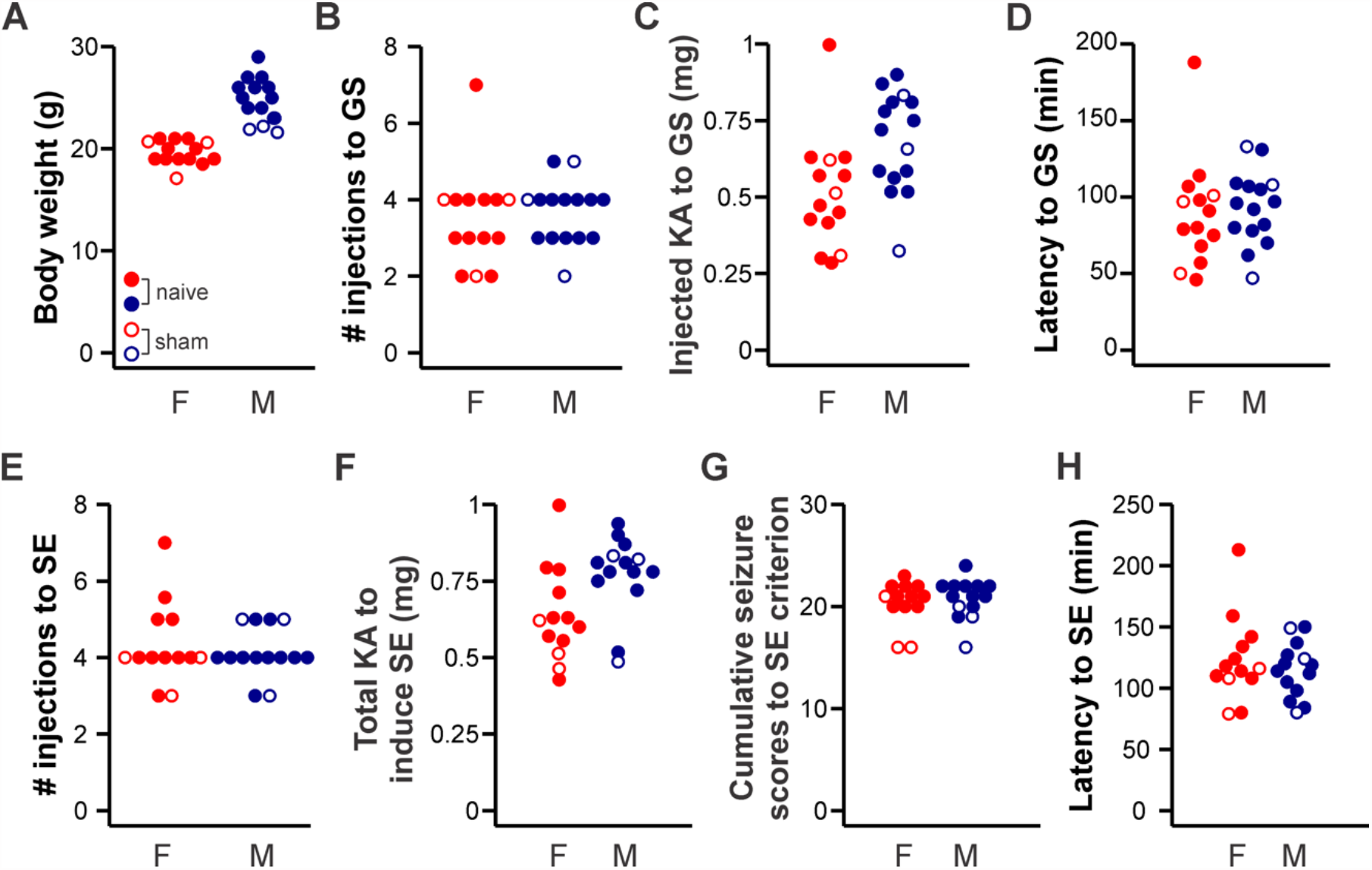
Similarity of data from naïve gonad-intact and sham surgery control mice. Individual values for naïve gonad-intact controls (filled circles) and gonad-intact controls that underwent sham surgical procedures (open circles). Females (F) = red, males (M) = blue.

## Notes

### Competing Interest Statement

The authors have declared no competing interest.

### Summary of Updates

Addition of new data and analyses, extensive edits to the text, addition of new references cited, addition of new supporting information section. The major conclusions of the previous version are unchanged.

